# Mitogenomics, phylogenetic position, and updated distribution of *Ergasilus kandti*, an ergasilid copepod parasitizing African cichlid fishes

**DOI:** 10.1101/2024.03.27.586948

**Authors:** Dries Jansen, Maarten P. M. Vanhove, Lawrence Makasa, Jiří Vorel, Nikol Kmentová, Armando J. Cruz-Laufer

**Author notes:** these authors contributed equally (Nikol Kmentová, Armando J. Cruz-Laufer). **Corresponding author:** Armando J. Cruz-Laufer.

## Abstract

Ergasilidae are a family of globally distributed copepods parasitizing freshwater fish. Despite their widespread occurrence, their phylogeographic patterns are poorly understood, specifically in the African Great Lakes. Here, we aim to provide an update on distribution of *Ergasilus kandti,* a copepod species infecting *Tylochromis polylepis,* an endemic cichlid fish species in Lake Tanganyika, and the phylogenetic relationship of African ergasilids. We present the first record of *E. kandti* parasitizing the gills of *T. polylepis* in Lake Tanganyika proper, identified through light microcopy and, for the first time for any ergasilid, confocal laser scanning microscopy. We suggest that this technique adds spatial context to characters and are hardly visible while using light microscopy. Phylogenetic analyses based on ribosomal DNA fragments suggest two monophyletic groups of African ergasilids. However, the phylogenetic relationships of *Ergasilus* remain unresolved, possibly because of the insufficient resolution of these widely used phylogenetic markers and low taxonomic coverage. A comparison of ergasilid mitochondrial genomes highlights traits found in other parasite lineages including genome shrinkage and low evolutionary rates of the *cox1* gene. This study presents the most extensive molecular characterization of any ergasilid species to date.

## Introduction

Copepods are among the most abundant multicellular organisms on Earth (Humes, 1994; Walter & Boxshall, 2023). Parasitic copepods with their free-living larval stages constitute a major proportion of this abundance. Among these, Ergasilidae Burmeister, 1835 is one of most species-rich taxa containing 275 described species infecting mostly teleost fishes (Walter & Boxshall, 2023). Ergasilid copepods pursue a unique life cycle among copepods: they complete their first developmental stages (nauplius to adult) as plankton and after mating, only fertilized females infest a host for the final parasitic phase of the life cycle (Boxshall & Defaye, 2008). As fish is a crucial component of the human food industry, understanding their parasites is key for sustainable food production. Copepods can be serious pests in fish, e.g. *Lernaea cyprinacea* Linnaeus, 1758 and *Ergasilus sieboldi* von Nordmann, 1832, both of which may cause substantial mortality and commercial losses in fish aquaculture (Boxshall & Defaye, 2008). This highlights the importance of studying copepods, and specifically parasitic copepods, yet, despite this relevance, the diversity of parasitic copepods remains understudied (Bernot et al., 2021).

Ancient lakes are biodiversity hotspots, including a high species richness of copepods. Together with Lake Baikal with over 120 species (Boxshall et al., 1993), Lake Tanganyika in East Africa is a prime example with 69 copepod species reported to date (including 34 endemic species) (Boxshall & Strong, 2006). Lake Tanganyika is the longest (660 km) and the second deepest (1436 m) lake in the world (O’Reilly et al., 2003) with an exceptional level of endemism (∼95%) of e.g. fishes, snails, and ostracods (Salzburger et al., 2014). These species-rich communities have often resulted from adaptive radiations, a rapid diversification event associated with adaptations to various available ecological niches (Schluter, 2000; Salzburger et al., 2014). The best-studied example of such a community in Lake Tanganyika are cichlid fishes (Cichlidae Bonaparte, 1835), which have become a model system for studying evolution and speciation (Salzburger et al., 2014; Salzburger, 2018; Ronco et al., 2020). Concerning parasitological research in Lake Tanganyika, the majority of studies focus on monogenean flatworm parasites infecting the gills of these cichlids (Vanhove et al., 2011; Rahmouni et al., 2017; Kmentová et al., 2021). Parasitic copepods occupy some of the same host organs as monogenean flatworms (e.g. fish gills) yet receive far less attention. Hitherto, eight species of parasitic copepods belonging to Ergasilidae have been reported in the lake, all of which belong to *Ergasilus* von Nordmann, 1832 (Míč et al., 2023). Notably, copepods dominate the lake’s zooplankton community whereas in most other lakes in East Africa cladoceran crustaceans are dominant, highlighting the importance of copepods to Lake Tanganyika’s nutrient cycle and ecosystem functioning (Sarvala et al., 1999). Given the combination of free-living and parasitic life stages in ergasilid copepods, knowledge on fish-copepod interactions, including parasitic relationships, are an essential component for lake-wide ecosystem dynamics. However, phylogeographic relationships of parasitic copepods are generally underexplored as only 3% of all copepod species and only seven out of 30 accepted genera of Ergasilidae have been examined in a molecular phylogenetic context (Bernot et al., 2021; Míč et al., 2023). Most of the molecular data on species of *Ergasilus* were published by Song et al. (2008), in a first phylogenetic analysis of the family with most specimens sampled in China. The first molecular data of African ergasilids have been released for five species of the eight so far reported in Lake Tanganyika (Míč et al., 2023) and one from Madagascar (Míč et al., 2024).

The cichlids of Lake Tanganyika all evolved from a common ancestor after the formation of the lake 9–12 Mya, except for two species: *Tylochromis polylepis* Boulenger, 1900 and *Oreochromis tanganicae* Günther, 1894. These last two colonized the lake only recently (510,000 years ago for *T. polylepis*), establishing themselves in an already mature adaptive radiation occupied by species with fine-tuned niche segregation (Koch et al., 2007). *Tylochromis polylepis* is less specialized and more of a generalist in ecological terms: The species is a bottom feeder, usually found above muddy and sandy bottoms close to the shore and river mouths (Koch et al., 2007) and feeds on gastropods, insect larvae, and plant material (Lamboj, 2004). As the lakes’ only tylochromine cichlid, *T. polylepis* contrasts with the many other cichlid lineages that have undergone adaptive radiation. Its parasites can, therefore, serve as a model of understanding how the absence of ecological differentiation of host species affects their parasite fauna. To date, only four parasite species including three monogenean flatworm species [*Cichlidogyrus mulimbwai* Muterezi Bukinga, Vanhove, Van Steenberge & Pariselle, 2012, *C. muzumanii* Muterezi Bukinga, Vanhove, Van Steenberge & Pariselle, 2012, and *C. sergemorandi* Rahmouni, Vanhove & Šimková, 2018] (Muterezi Bukinga et al., 2012; Rahmouni et al., 2018) and a single copepod species (*Ergasilus kandti* van Douwe, 1912) have been reported to infect *T. polylepis*.

This study investigates the copepod *E. kandti*, a widely distributed species (Fig. 1, Supplementary Table S1) infecting a specimen of *T. polylepis* in Lake Tanganyika. The species has previously been reported from Lake Tanganyika parasitizing *Limnotilapia dardennii* Boulenger, 1899, *Lamprologus lemairii* Boulenger, 1899, *Plecodus paradoxus* Boulenger, 1898, *Pseudosimochromis curvifrons* Poll, 1942, *Oreochromis* sp. (cited as *Tilapia* sp.), and *O. tanganicae* (Capart, 1944; Fryer, 1965). Furthermore, *E. kandti* has also been reported from *T. polylepis* in the Malagarasi Delta (Tanzania) that borders the lake (Fryer, 1967). Therefore, our study provides the first record of *E. kandti* on *T. polylepis* in Lake Tanganyika proper. The species has also been reported infecting *Lates niloticu*s (Linnaeus, 1758) and *Bagrus bajad* (Fabricius, 1775) in Lake Albert (Fryer, 1965; Thurston, 1970), *Pterochromis congicus* (Boulenger, 1897) in Lake Tumba (Fryer, 1964), *Tylochromis bangwelensis* Regan, 1920 and *T. mylodon* Regan, 1920 in Lake Mweru (Fryer, 1967), *Citharinus citharus* (Geoffroy Saint-Hilaire, 1808), *Hemisynodontis* sp., *L. niloticus*, and *Synodontis membranaceus* (Geoffroy Saint-Hilaire, 1809) in the Black and White Volta confluence (Ghana) (Paperna, 1969), and *L. niloticus* in the River Niger (Capart, 1956; Fryer, 1968) and the Lower Nile (Fryer, 1968). Here, we provide a morphological comparison with other ergasilid species through light and confocal laser microscopy, the latter of which is applied to ergasilids for the first time. We conduct phylogenetic analyses of all species of Ergasilidae of which molecular data are currently available and assess the relationship of African representatives. We also provide the first ergasilid whole-genome sequencing reads [but see previous studies employing Sanger sequencing to obtain complete mitochondrial genome sequences, see Feng et al. (2016), He et al. (2023), and Hua et al. (2023)], the first complete mitogenome sequence and ribosomal operon of any African ergasilid species, and the first quantitative comparison of mitochondrial genome features of ergasilid copepods.

**Fig. 1.**
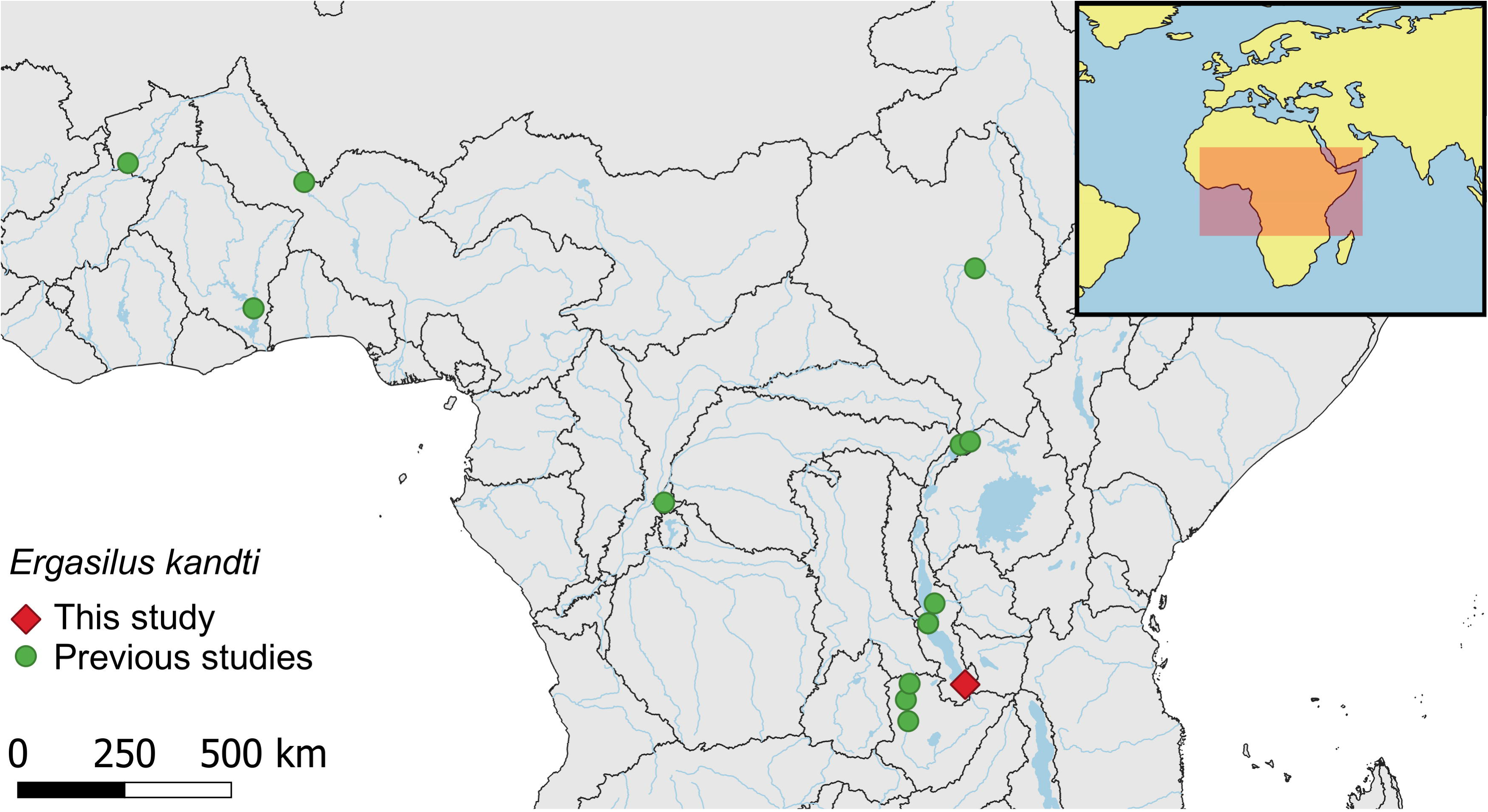
Recorded locations of *Ergasilus kandti*. Geographical coordinates were estimated based on reports in the literature (i.e. geographical centers of lakes, approximate locations indicated in figures etc.) (see Supplementary Table S1). Borders indicate limits of freshwater ecoregions according to Thieme et al. (2005).

## Materials and methods

### Morphological analysis

Copepod specimens (n = 23) were collected from the gills of the single specimen of *T. polylepis* purchased from a fisherman at the market in Mpulungu, Zambia (−08.767, 031.117) on September 20th, 2019. Specimens were stored in 70% ethanol at 4°C. The samples were stained with Congo Red (1 mg/ml aqueous solution) for a minimum of 24 h at room temperature. The dye proved to be stable compared to other fluorescence dyes (Michels & Büntzow, 2010). Following the staining procedure, the samples were rinsed in deionized water for 30 min. The specimens were mounted on concave microscope slides (to prevent crushing by the cover clip) in glycerin. Copepod specimens were imaged with a confocal laser microscope (ZEISS LSM900) with an excitation of 561 nm, an emission at 570–670 nm, pinhole size of 1 AU (airy unit); laser intensity of 0,2%, 400x magnification; resolution of 512 x 512 px, bidirectional scanning, eight-fold averaging, 30 tiles, and Z-stack with interval of 1–1.12 µm. Tile images were stitched with the software ZEISS ZEN version 3.1 (Carl Zeiss, Jena, Germany). Images were further processed in Fiji/ImageJ v1.54d (Schindelin et al., 2012). To adjust the contrast of dimmed regions of the image, the *StackContrastAdjustment* plugin was used (Čapek et al., 2006), followed by maximum-intensity projections for two-dimensional representation of the images and denoising. Specimens were identified to species level using a brightfield light microscope (Leica DM2500) based on the spine-seta formulae provided by (Capart, 1944) and additional characteristics listed in previous studies (Oldewage & van As, 1988; Schlebusch, 2014).

### DNA extraction and PCRs

DNA extractions were performed in a UV cabinet and all utensils were sterilized and UV–irradiated before usage. The extractions were completed with a slightly modified version of Kmentová et al. (2021), applying an extended enzymatic digestion period (55°C, 400 rpm, three hours) to digest the soft tissues of the specimen. Exoskeletons were removed after this digestion step. The exoskeletons were stained with Congo Red as mentioned above but mounted in glycerin on standard (non-concave) microscopic slides. For permanent storage, specimens were either mounted in glycerin in sealed 1.5-ml tubes or in Euparal on microscopic slides. Voucher material was deposited in the collection of Hasselt University under accession numbers HU XXIII.1.41–HU XXIII.1.50, HU XXIII.2.01, and HU T-I.1–HU T-I.11.

The PCR reaction targeting a portion of the 28S rDNA region contained forward primer 28SF (5’ - ACA ACT GTG ATG CCC TTA - 3’) and reverse primer 28SR (5’ - TGG TCC GTG TTT CAA GAC - 3’) (Song et al., 2008). The PCR targeting a portion of the 18S rDNA region contained forward primer 18SF (5’ - AAG GTG TGM CCT ATC AAC - 3’) and reverse primer 18SR (5’- TTA CTT CCT CTA AAC GCT - 3’) (Song et al., 2008). An identical master mix was used for both PCRs. It contained 1x PCR buffer, 0.2mM dNTPs, 0.5µM of each primer, 0.5U of Q5 polymerase, 2µl of DNA and ddH2O was added to bring the total volume to 25µl. The PCR conditions were set as follows: 5 min at 94°C; 35 cycles of 30 s at 94°C, 1 min at 54°C, 1 min at 72°C; a final elongation of 10 min at 72°C. Gel electrophoresis was used to check the PCR products for successful amplification. The PCR products were purified using a GeneJET PCR Purification Kit (Thermo Fisher Scientific, Waltham, MA, USA) and Sanger sequencing was outsourced to Macrogen Europe (Amsterdam, The Netherlands). Sequence quality was checked visually through the sequencing chromatograms. The 28S rDNA and 18S rDNA sequences were uploaded to GenBank and can be found under accession numbers PQ249839 and PQ249841X–PQ249843, respectively. We also attempted the amplification of a fragment of the cytochrome *c* oxidase subunit I gene (*cox1*) using a set of general invertebrate primers (Folmer et al., 1994; Nadler et al., 2006) previously applied successfully to ergasilids (Santacruz et al., 2020; Fikiye et al., 2023). However, none of these reactions were successful at the amplification stage, possibly due to age of the samples and the low primer specificity.

### Whole-genome sequencing and assembly of ribosomal operon and mitogenome

The extracted genomic DNA of a single copepod individual was subjected to the generation of a genomic read pool. Library preparation and sequencing on the Illumina NovaSeq6000 platform (short-insert paired-end sequencing, 2x 150 bp configuration) were outsourced to Novogene Europe (Cambridge, UK). For the assembly of the ribosomal operon, raw reads were trimmed through Trimmomatic v0.39 (Bolger et al., 2014) using a sliding window approach (settings: *SLIDINGWINDOW:4:28 HEADCROP:5 MINLEN:100 ILLUMINACLIP:TruSeq3-PE.fa:2:30:10:2:True*). The quality of filtered reads was checked in FastQC v0.11.8 (Andrews, 2018). Reads of the ribosomal operon were baited through complete sequences of the full ribosomal operon of other copepod species (Ki et al., 2011; Weydmann et al., 2017) using *mirabait* in Mira V5rc1 (Chevreux B & Suhai, 1999). The operon was assembled *de novo* through Spades v3.15.2 (Prjibelski et al., 2020) with a range of Kmer lengths (21, 33, 55, 77, 99, and 127). The resulting assembly graphs were visualized in Bandage v0.8.1 (Wick et al., 2015). The resulting contigs were subjected to a BLAST search on NCBI GenBank to identify those belonging to a copepod. The selected contig sequence was then realigned to the bait sequences (see above) using MAFFT v.7.409 to locate the boundaries of the rDNA and transcribed spacer regions.

The mitogenome of *E. kandti* was assembled using raw reads and part of the cytochrome *c* oxidase subunit I mitochondrial DNA (*cox1*) of *Ergasilus* sp. (GenBank accession number KU557411) as a seed in NOVOPlasty v4.3.1 with a k-mer length of 21, read length of 151 and insert size of 300. Results from the MITOS web server (code: invertebrate mitochondrial) (Bernt et al., 2013) were used for mitogenome annotation combined with the tRNAscan-SE (Lowe & Chan, 2016) and RNAfold (Lorenz et al., 2011) web servers to verify the tRNAcoding regions. The annotation was verified via a visualization of open reading frames and aligned with available ergasilid mitogenomes [*Ergasilus tumidus* Markevich, 1940 - OQ596537, NC_073502; *Sinergasilus major* (Markevich, 1940) – OQ160840, NC_081952; *Sinergasilus polycolpus* (Markevich, 1940) – KR263117, EU621723, NC_028085; *Sinergasilus undulatus* (Markevich, 1940) – MW080644, NC_054173] in Geneious Prime 2023.2.1 (Auckland, New Zealand). The annotated mitochondrial genome of *E. kandti* was visualized using CGView (Grant et al., 2023) including the information on GC content and GC skew+/-. Protein coding rDNA regions were aligned separately using Clustal Omega v 1.2.2 and subsequently concatenated in Geneious Prime 2023.2.1. The sliding window analysis and dN/dS ratio across all 13 PCRs was inferred through DnaSP v5 (Librado & Rozas, 2009) to summarize nucleotide polymorphisms across all ergasilid copepod species of which the mitochondrial genomes were assembled to date (see above). The assembled nuclear ribosomal and mitogenome DNA sequences of *E. kandti* are available in the GenBank Nucleotide Database under the accession numbers PQ249840 and PQ276880. Raw Illumina reads were submitted to SRA (accession number: SRR30471034) under BioProject accession PRJNA1153390.

### Phylogenetic reconstruction

To infer the evolutionary history of *E. kandti*, we assembled multiple sequence alignments of partial 18S and 28S nuclear ribosomal DNA (18S and 28S rDNA) and *cox1* combining sequences generated during this study with sequences accessed on NCBI GenBank (Table 1). Due to the close relationship of species belonging to Lernaeidae and Ergasilidae (Song et al., 2008; Kvach et al., 2021; Abdel-Radi et al., 2022), three representatives of Lernaeidae were chosen as outgroups. Sequences of three species of Mytilicolidae Bocquet & Stock, 1957 and two species of Myicolidae Yamaguti, 1936 were also added to the alignment because previous studies indicate close ties of these families with Ergasilidae (Khodami et al., 2019; Oliveira et al., 2021). We aligned 18S and 28S rDNA sequences using *L-INS-i* in MAFFT v.7.409 (Katoh & Standley, 2013) and removed poorly aligned positions and divergent regions using the options for less stringent parameters (options: - b2= <*minimum*> -b3=10 -b4=5 -b5=h) in Gblocks v.0.91b (Talavera & Castresana, 2007). For *cox1*, sequences were aligned in MACSE v2.06 (Ranwez et al., 2011, 2018) using the algorithm *trimNonHomologousFragments*, *alignSequences*, and *TrimSequenceAlignment* under default parameters and with the Invertebrate Mitochondrial Code.

**Table 1:**
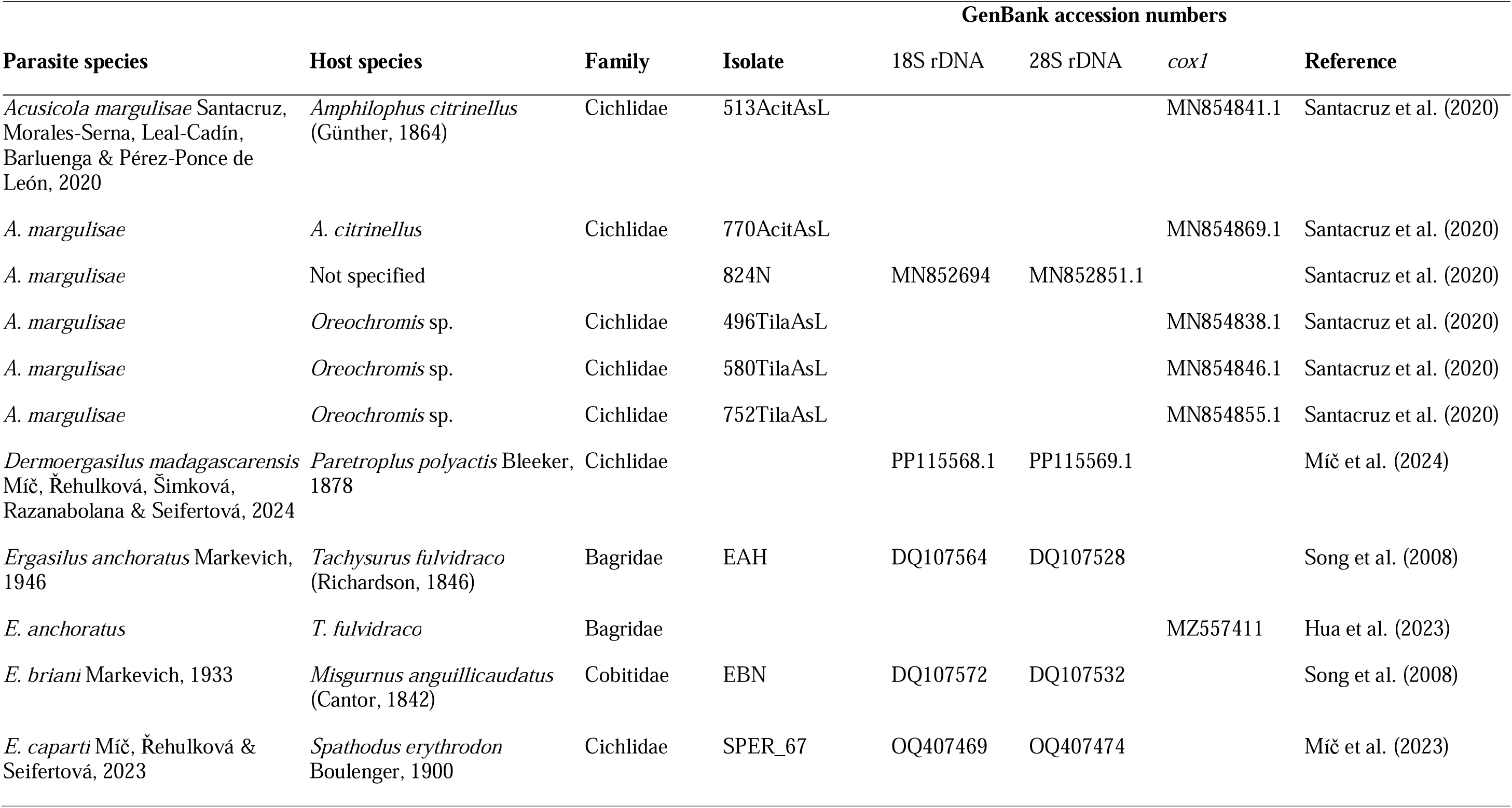
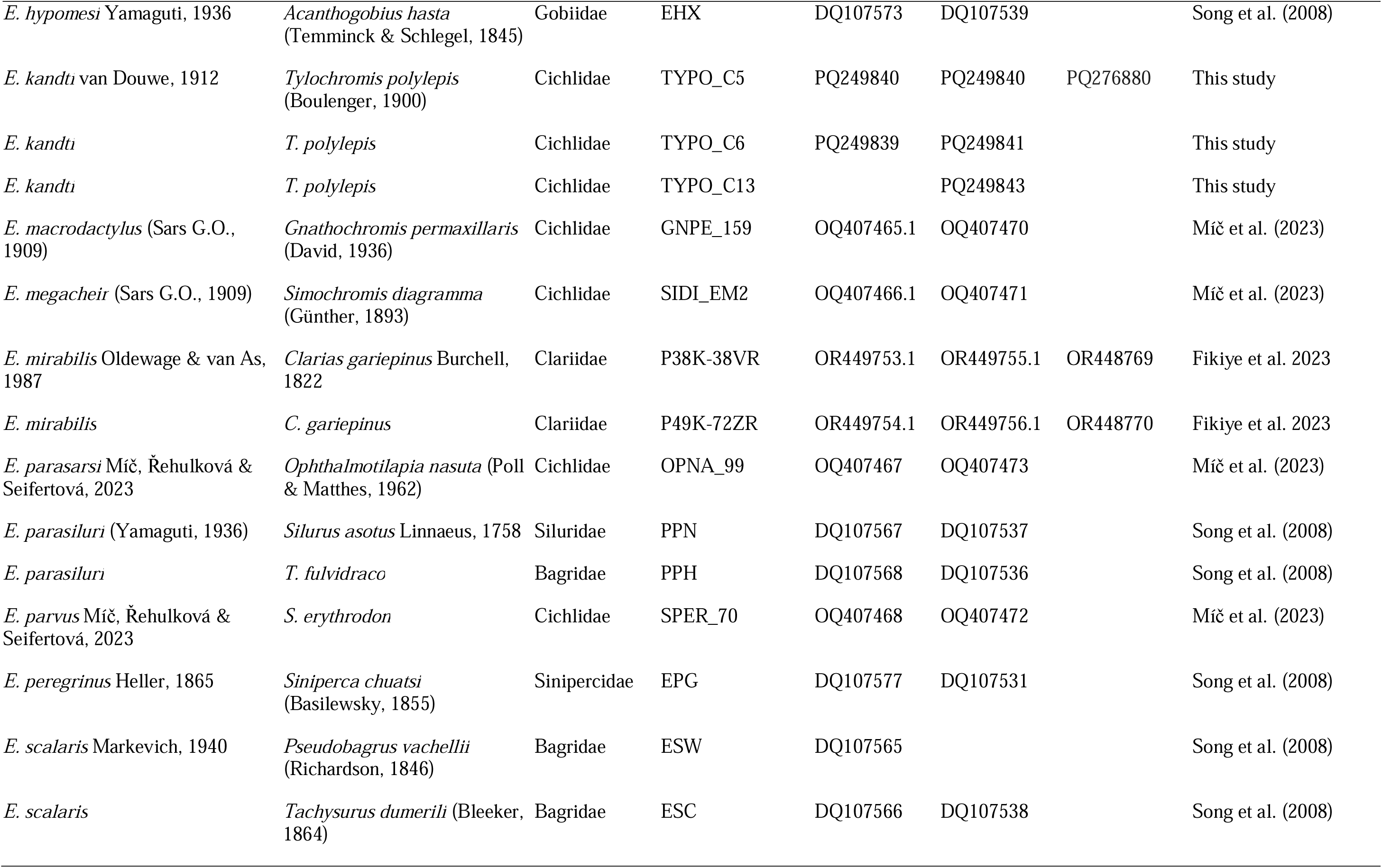
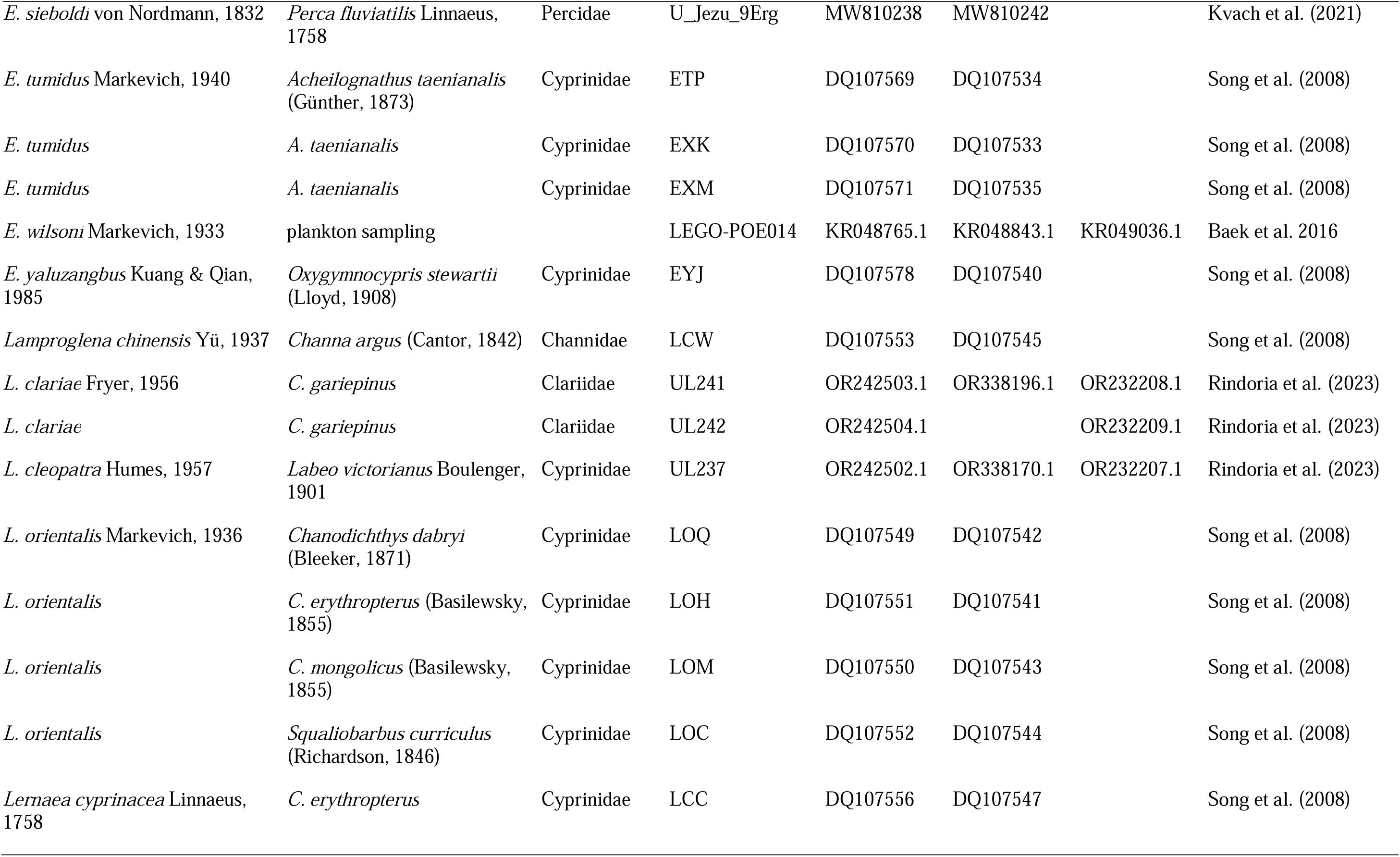
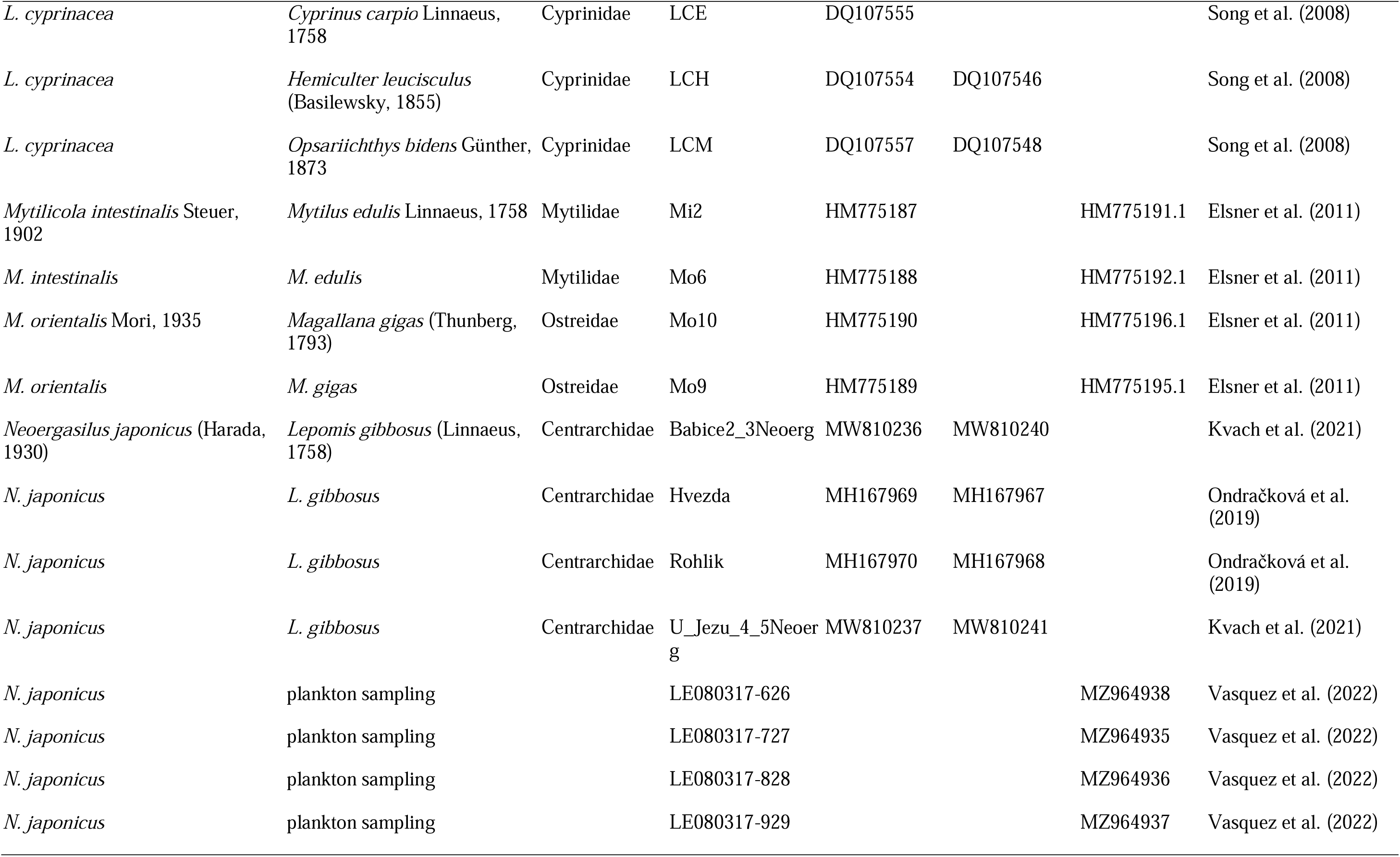
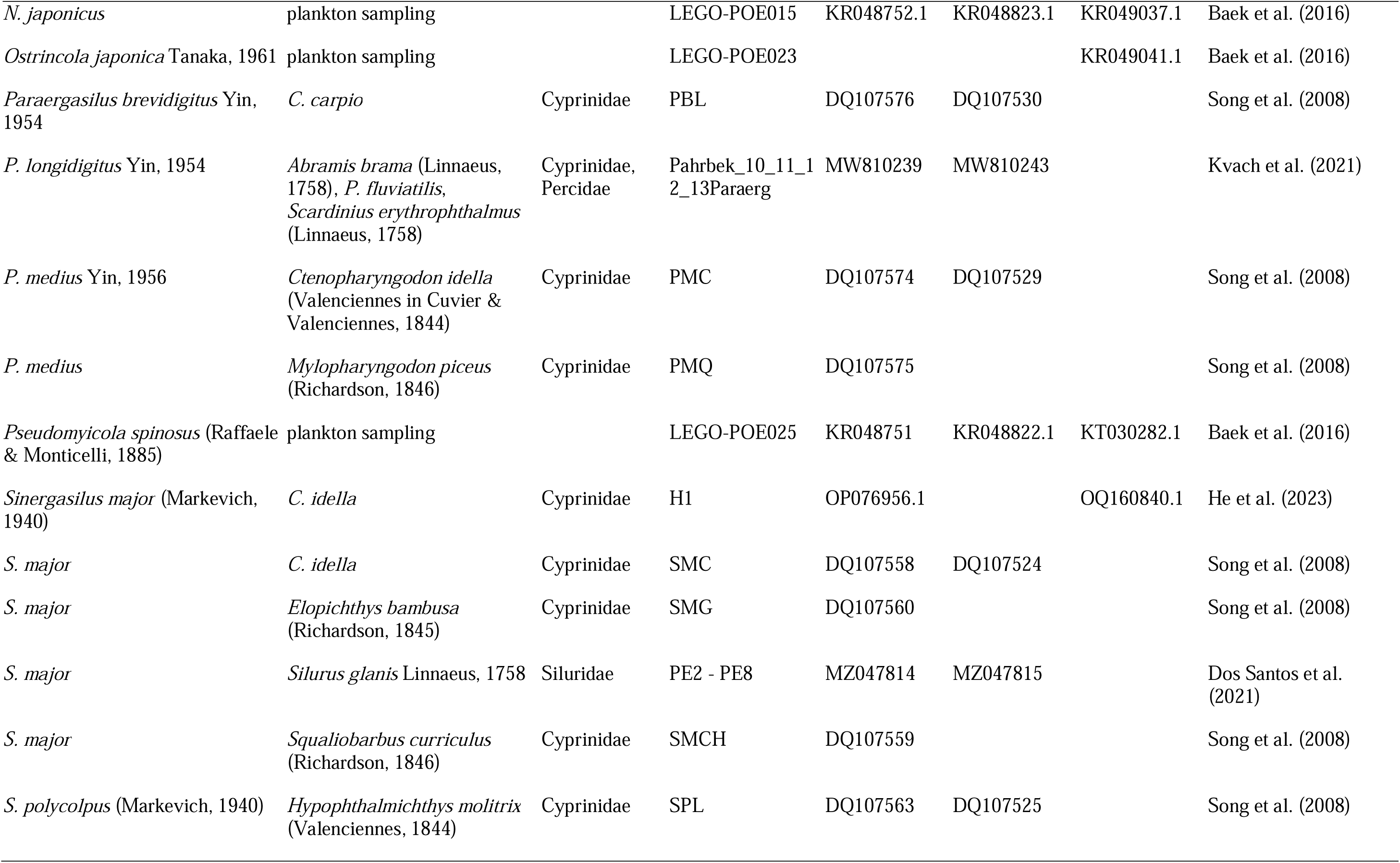
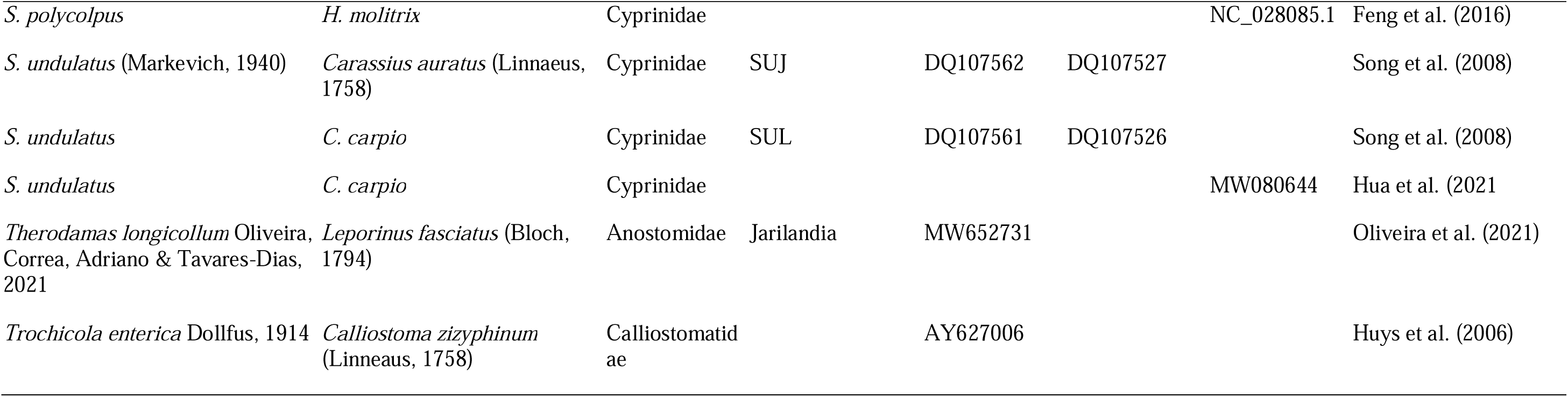
Specimen data of copepods used for phylogenetic analyses including host species, GenBank accession numbers, and reference study.

We estimated tree topologies through maximum likelihood (ML), Bayesian inference (BI), and maximum parsimony (MP) methods using IQ-Tree v2.2.26 (Minh et al., 2020), MrBayes v3.2.7 (Ronquist et al., 2012), and TNT v1.6 (Goloboff et al., 2008; Goloboff & Morales, 2023), respectively. Because of the minimal overlap in terms of species and specimens between the nuclear and mitochondrial loci, we opted to concatenate only the 18S and 28S rDNA and analyze *cox1* separately. The alignments were partitioned by locus and, for *cox1*, by codon position. For ML and BI, we selected the substitution models for each partition according to the Bayesian information criterion (BIC) as suggested by ModelFinder in IQ-Tree (Kalyaanamoorthy et al., 2017) using partition merging (Chernomor et al., 2016) (Table 2). For ML analyses, we estimated branch support values using both ultrafast bootstrap approximation (UF-Boot) (Hoang et al., 2018) and Shimodaira–Hasegawa-like approximate likelihood ratio tests (SH-aLRT) (Guindon et al., 2010) with 10,000 replicates. To minimise false positives, UF-Boot was estimated by resampling partitions as well as sites within resampled partitions (Gadagkar et al., 2005; Seo et al., 2005). For BI analyses, model selection in ModelFinder was restricted to models implemented in MrBayes (Table 2). We used two runs and four chains of Metropolis-coupled Markov chain Monte Carlo (MCMC) iterations with calculation conducted in parallel to optimize runtime (Altekar et al., 2004). We ran 20 million generations with a burn-in fraction of 0.25 and sampled the trees every 1000th generation. We checked convergence criteria by assessing the average standard deviation of split frequencies (<0.01 in all datasets) and the effective sample size (>200) using Tracer v1.7 (Rambaut et al., 2018).

**Table 2:** Best-fitting nucleotide substitution models for alignments of 18S rDNA, 28S rDNA, and each codon position of COI mDNA selected by ModelFinder for use in IQ-Tree and MrBayes.

| Alignment | Base pairs | Model IQ-Tree | Model MrBayes |
| --- | --- | --- | --- |
| 18S rDNA | 1500 | TIM3e+I+R2 | SYM+I+G4 |
| 28S rDNA | 754 | GTR+F+I+G4 | GTR+F+I+G4 |
| *Cox1* | 1551 | TIM2+F+G4 | GTR+F+G4 |
|  |  | K3Pu+F+G4 | F81+F+G4 |
|  |  | HKY+F+I+R2 | HKY+F+G4 |

For analyses in TNT, we used extended implied weighting (command *xpiwe*) (Goloboff, 2014) for different values of the concavity constant k (20, 21, 23, 26, 30, 35, 41, 48, 56). Extended implied weighting minimizes the impact of characters with missing data that are weighted artificially high in the original implied weighting method (Goloboff, 1993). For the rDNA data, we explored two weighting schemes proposed by Mirande (2019): all characters weighted separately (SEP) and all characters weighted according to the average homoplasy of the marker (BLK). For *cox1*, we tested an additional scheme: characters weighted according to the average homoplasy of their codon position (POS). Similar to Mirande (2009), we selected the k-value and weighting scheme that produced the most stable topology to compute a strict consensus tree. The distortion coefficient and subtree pruning and regrafting (SPR) distance were used as selection criteria by computing the similarity of each consensus tree to the others obtained under the different parameters (commands *tcomp* and *sprdiff*). MP tree searches involved rounds of tree fusing, sectorial searches, and tree drifting (Goloboff, 1999) under default settings and each round was stopped following three hits of the same optimum. Gaps were treated as missing data. As bootstrapping and jackknifing are reported to be distorted by differently weighted characters (Goloboff, 2003), branch support was estimated through symmetrical resampling (probability of change of 0.33) and values expressed as differences in frequencies (GC, “Groups present/Contradicted”). We considered a BI posterior probability ≥ 0.95, UF-Boot ≥ 95 and SH-aLRT statistic ≥ 80 (Hoang et al., 2018), and GC ≥ 0 as well-supported. Nodes without support above these thresholds in any of the phylogenetic trees were collapsed into polytomies for visualization. Trees were visualized through R v4.3.2 (R Core Team, 2023) packages *ggplot2* v3.4.4 (Wickham, 2016) and *ggtree* v3.10.0 (Yu et al., 2017, 2018).

## Results

### Imaging and species identification

The collected specimens were identified as *E. kandti* based on characters observed through light and laser confocal microscopy (Fig. 2, 3). Congo red showed no signs of fading or bleaching several months after the initial staining and the multiple imaging sessions of up to 12 hours. All specimens presented characters associated with the genus *Ergasilus*, i.e. a short, drop-shaped body (Fig. 2a, 3a, 3c) with a sclerotized terminal segment of the second antennae and a single point (Kuchta et al., 2018). Furthermore, the specimens all presented a two-segmented exopod on the fourth leg (in contrast with the rest of the legs that all have three segmented exopods), which is also characteristic of species of *Ergasilus* (see Fryer, 1965). All of the specimens examined in detail (n = 10) presented a characteristic combination of the following morphological features specific to *E. kandti*: a fold/groove shaped like an inverted T on the carapace posterior to the eyespot (Fig. 3a), and a circular cephalic structure anterior to the fold/groove (Fig. 3a), a pair of fifth legs reduced to a single segment with four setae (Fig. 3b), a pair of modified large grasper-like antennae with a tooth-like projection on the distal inner side of the third segment (Fig. 2b, 3d), a pair of five-segmented antennules with four setae on the distal segments (Fig. 2c and 3e), a pair of caudal rami that terminates in two long setae and two short setae (Fig. 2d). Specifically, the tooth-like projection associated with the large grasper-like antennae has been suggested to be a unique character of the species among species of *Ergasilus* from Lake Tanganyika and the rest of Africa (Oldewage & van As, 1988; Míč et al., 2023).

**Fig. 2.**
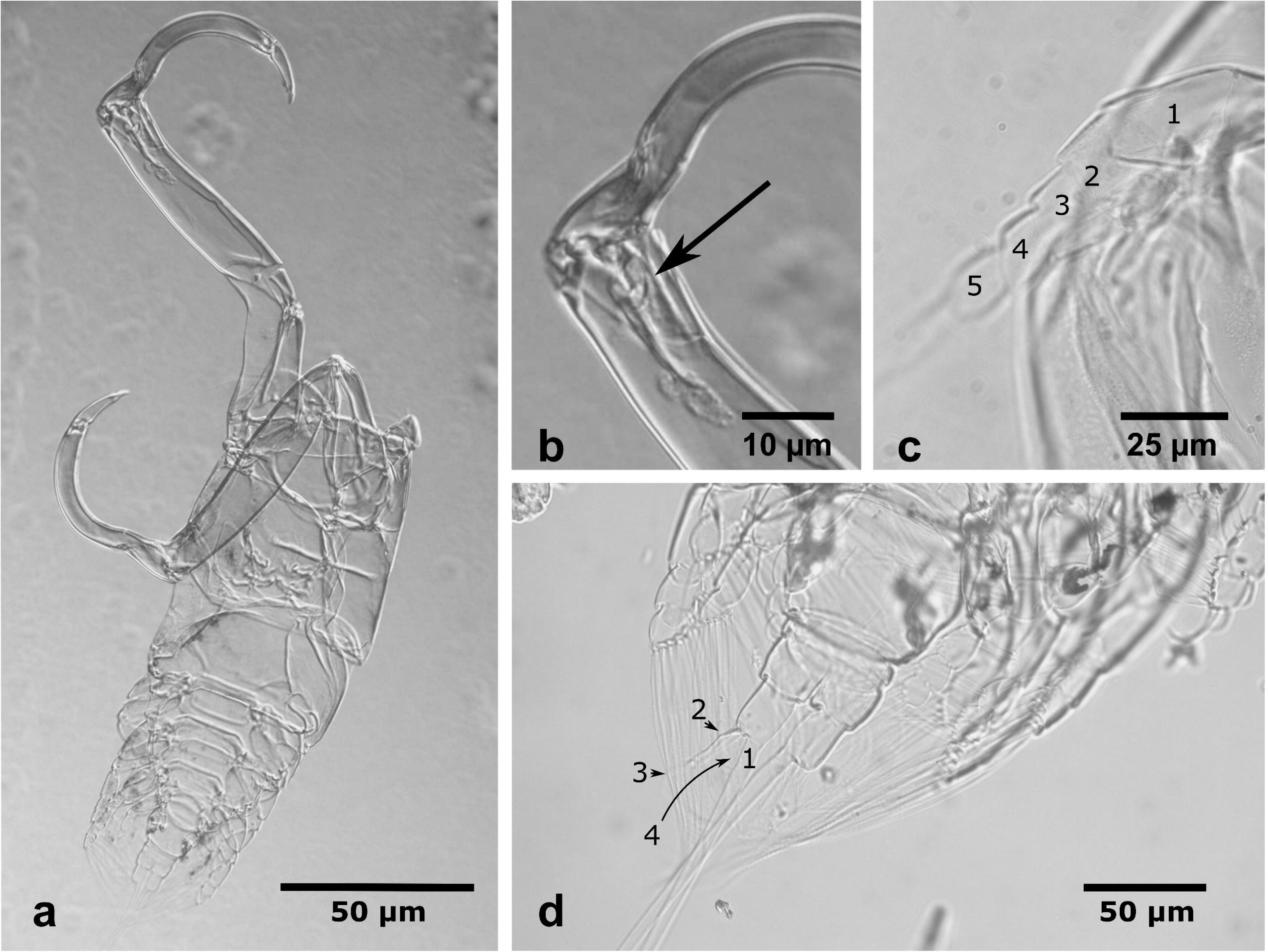
Light microscope images of *Ergasilus kandti*. a: whole specimen, ventral view. b: detail tooth-like projection on third segment of second antenna. c: detail of the five-segmented antennule. d: caudal rami that terminates in two long (1, 3) and two short (2, 4) setae.

**Fig. 3:**
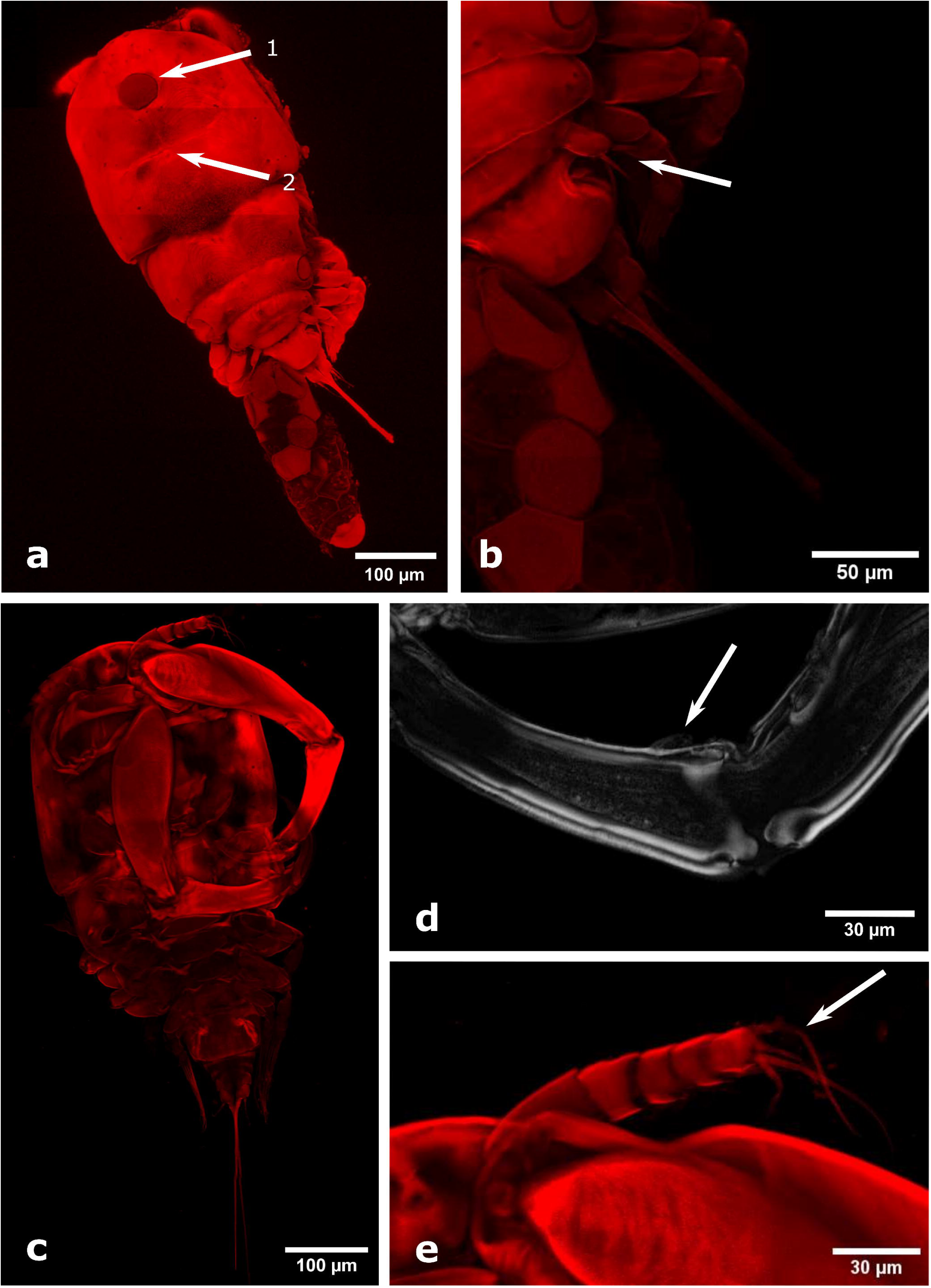
Confocal laser microscopic images of *Ergasilus kandti*. a: dorsal view with (1) eyespot (circular cephalic structure) and (2) characteristic groove in the shape of an inverse T. b: detail of the reduced fifth leg, characteristic of species of *Ergasilus*. c: ventral view. d: detail of second antenna with lightly stained tooth-like projection (see arrow, greyscale was used for better contrast). e: detail of the five-segmented antennule with four setae at distal end.

Concerning the spine-seta formula, seven out of ten specimens were identical to the original description (Capart, 1944). Occasionally, deviations in the numbers of setae were observed, i.e. on the second segment of the endopodite of the third leg for a single specimen (see Supplementary Table S2) compared to two setae mentioned in the original description. However, these likely broke off during the mounting process as all species of *Ergasilus* have one or two setae on this leg segment. Apart from the abovementioned characters, African congeneric species were ruled out through the comparison of their respective spine-seta formulas. *Ergasilus cunningtoni* Capart, 1944 and *E. macrodactylus* (Sars, 1909) have four (Sars, 1909; Capart, 1944) and *E. nodosus* Wilson, 1924 five setae on the third segment of the exopodite of the third leg (Wilson, 1924) compared to the six setae present in *E. kandti* (Fig. 2b) (van Douwe, 1912). Furthermore, *E. cunningtoni, E. lamellifer* Fryer, 1961, *E. latus* Fryer, 1960, *E. macrodactylus, E. megacheir* Sars, 1909, *E. mirabilis* Oldewage & van As, 1987, *E. caparti* Míč, Řehulková & Seifertová, 2023, *E. parvus* Míč, Řehulková & Seifertová, 2023, *E. parasarsi* Míč, Řehulková & Seifertová, 2023, and *E. sarsi* Capart, 1944 have a six-segmented antennule compared to a five-segmented one present in *E. kandti* (Fig. 2a, 3d). *Ergasilus kandti* differs from *E. flaccidus* Fryer, 1965 and *E. nodosus* by having a single segmented fifth leg, and from *E. inflatipes* Cressey, 1970 by not having plumose setae on the fifth leg.

### DNA sequence assemblies

Through PCRs and Sanger sequencing, we obtained overall three rDNA sequences of three specimens of *E. kandti* (Table 1), the first from this species. Genomic DNA sequencing yielded 38,115,846 indexed paired-end reads. The complete ribosomal operon is 3,616 bp long (56.6% GC). Overall, for 28S rDNA (629–3838 bp) and 18S rDNA (1047–1849 bp), the sequences showed 99.8–99.9% similarity (uncorrected distance) for both markers. The complete mitochondrial genome of *E. kandti* is 14,639 bp long (35.8% GC) and includes 13 intron-free protein coding genes (PCGs), 22 tRNA genes, and two genes coding for the large and small subunits of the mitochondrial rRNA (Fig. 4, Table 3). The usage of T- and TA-abbreviated stop codon in the PCGs are three and two out of 13 PCGs, respectively. A comparison of available mitogenomes with other representatives of Ergasilidae highlighted differences concerning the gene order. *Ergasilus kandti* differs in the mutual position of *trnS1* and *trnS2* with *E. tumidus* (GenBank accession number NC_073502). Furthermore, we found that the mutual position of multiple tRNAs (*trnR, trnI, trnA* and *trnH*) differed between *S. polycolpus* and the four other ergasilid species, of which the mitochondrial genome sequences are available (*trnC, trnI, trnA, trnH, trnR*, and *trnT* in *S polycolpus*, and *trnC, trnR, trnI, trnA trnH*, and *trnT* in the rest). The existence of a control region or AT-loop region between *trnT* and *rrnL* was suggested by Hua et al. (2021) and Feng et al. (2016) for *S. majo*r (597 bp) and *S. polycolpus* (156 bp), respectively. In *E. kandti*, a non-coding region of 443 bp between *trnT* and *rrnL* (see Fig. 4) resembles this putative control region. The AT content in this non-coding region of *E. kandti* is 59.4% compared to 52.6% in *S. major* and 81.4% in *S. polycolpus*. We also observed that the length of rrnL in *S. polycolpus* (Feng et al., 2016) appears to be shorter compared to the other species including *E. kandti* (668 bp *vs*. 878 bp).

**Fig. 4:**
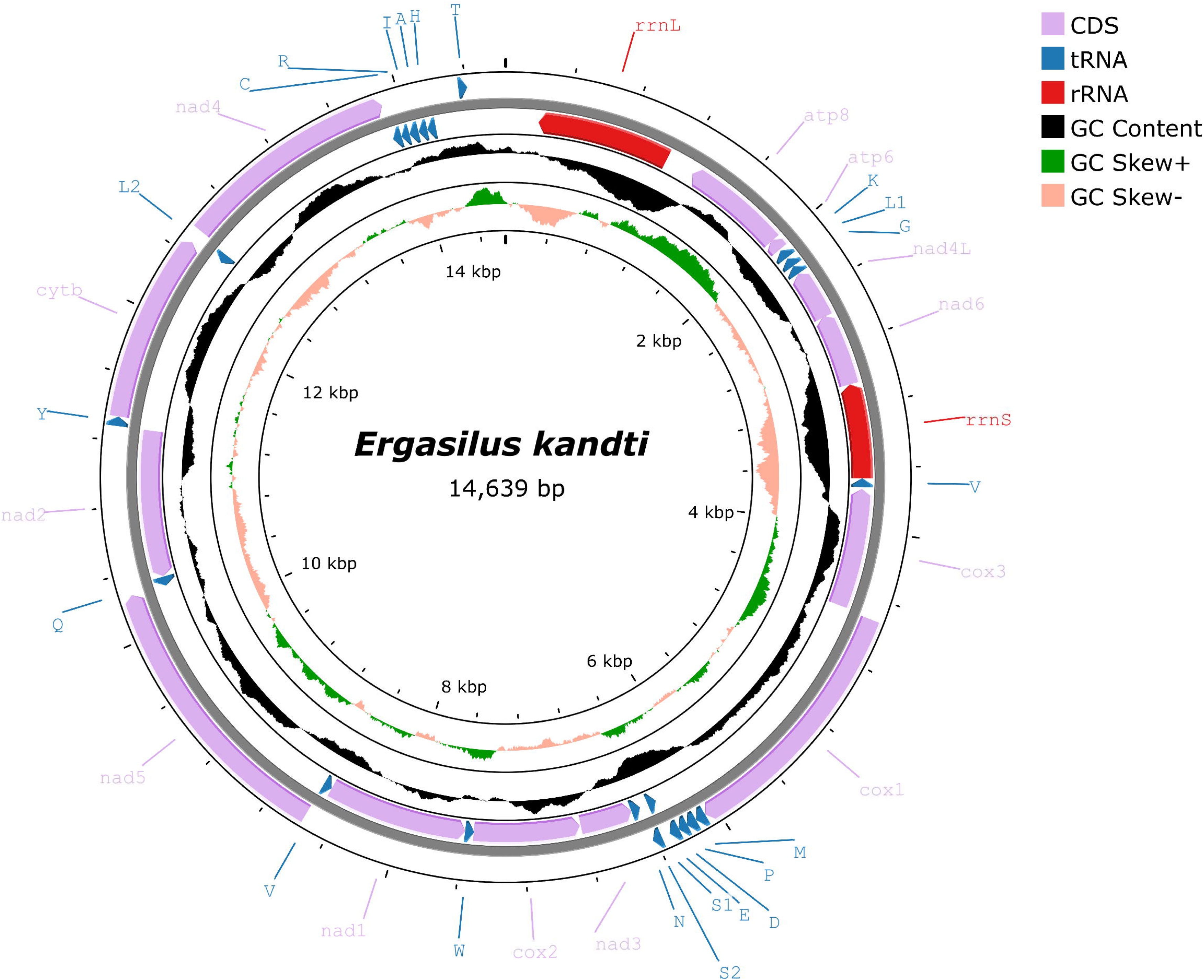
Mitochondrial genome of *Ergasilus kandti* with GC content (in black), positive (in green) and negative GC Skew (light orange) values.

**Table 3:** Annotation table of the mitochondrial genome of *Ergasilus kandti*.

| Region | Position | Length<br>(bp) | Start/stop codon | Anticodon |
| --- | --- | --- | --- | --- |
| **Non-coding region** | 1 -> 215, 14,412 -> 14,639 | 443 |  |  |
| **16S** | 1,093 -> 216 | 878 |  |  |
| **atp6** | 1,968 -> 1,285 | 684 | ATG/TAG |  |
| **atp8** | 2,044 -> 1,961 | 84 | ATA/TAG |  |
| **trnK** | 2,115 -> 2,054 | 62 |  | TTT |
| **trnL1** | 2,243 -> 2,185 | 59 |  | CAG |
| **trnG** | 2,243 -> 2,185 | 59 |  | TCC |
| **nad4L** | 2,574 -> 2,243 | 332 | ATG/TA |  |
| **nad6** | 3,046 -> 2,567 | 480 | ATA/TAG |  |
| **12S** | 3,667 -> 3,047 | 621 |  |  |
| **trnV** | 3,724 -> 3,667 | 58 |  | TAC |
| **cox3** | 4,514 -> 3,737 | 780 | ATG/T |  |
| **cox1** | 4,527 -> 6,075 | 1549 | ATA/T |  |
| **trnM** | 6,077 -> 6,139 | 63 |  | CAT |
| **trnP** | 6,140 -> 6,202 | 63 |  | TGG |
| **trnD** | 6,199 -> 6,256 | 58 |  | GTC |
| **trnE** | 6,255 -> 6,316 | 63 |  | TTC |
| **trnS1** | 6,366 -> 6,309 | 58 |  | TCT |
| **trnS2** | 6,367 -> 6,424 | 58 |  | AGT |
| **trnN** | 6,482 -> 6,421 | 62 |  | GTT |
| **nad3** | 6,826 -> 6,483 | 339 | ATA/TA |  |
| **cox2** | 7,529 -> 6,833 | 697 | TTG/T |  |
| **trnW** | 7,586 -> 7,527 | 60 |  | TCA |
| **nad1** | 7,529 -> 6,833 | 924 | ATA/TAA |  |
| **trnF** | 8,566 -> 8,509 | 58 |  | GAA |
| **nad5** | 8,568 -> 10,268 | 1701 | ATA/TAA |  |
| **trnQ** | 10,322 -> 10,261 | 62 |  | TTG |
| **nad2** | 14,358 -> 14,411 | 960 | ATA/TAA |  |
| **trnY** | 11,286 -> 11,343 | 58 |  | GTA |
| **cytb** | 11,344 -> 12,480 | 1137 | ATC/TAA |  |
| **trnL2** | 12,545 -> 12,485 | 61 |  | TAA |
| **nad4** | 12,564 -> 13,892 | 1329 | ATA/TAG |  |
| **trnC** | 13,948 -> 13,891 | 58 |  | GCA |
| **trnR** | 14,002 -> 13,948 | 55 |  | TCG |
| **trnI** | 14,063 -> 14,003 | 61 |  | GAT |
| **trnA** | 14,122 -> 14,065 | 58 |  | TGC |
| **trnH** | 14,180 -> 14,123 | 58 |  | GTG |
| **trnT** | 14,358 -> 14411 | 54 |  | ACA |

The sliding window analysis indicated an overall higher nucleotide diversity in genes coding for dehydrogenase subunits compared to other protein coding regions of cytochromes and ATP synthases (see Fig. 5). The lowest values were found for the *cox1* and *cox2* genes, respectively. The dN/dS ratios follow a similar trend with the highest values reported for nad6 and the lowest for *cox1* (see Fig. 6).

**Fig. 5:**
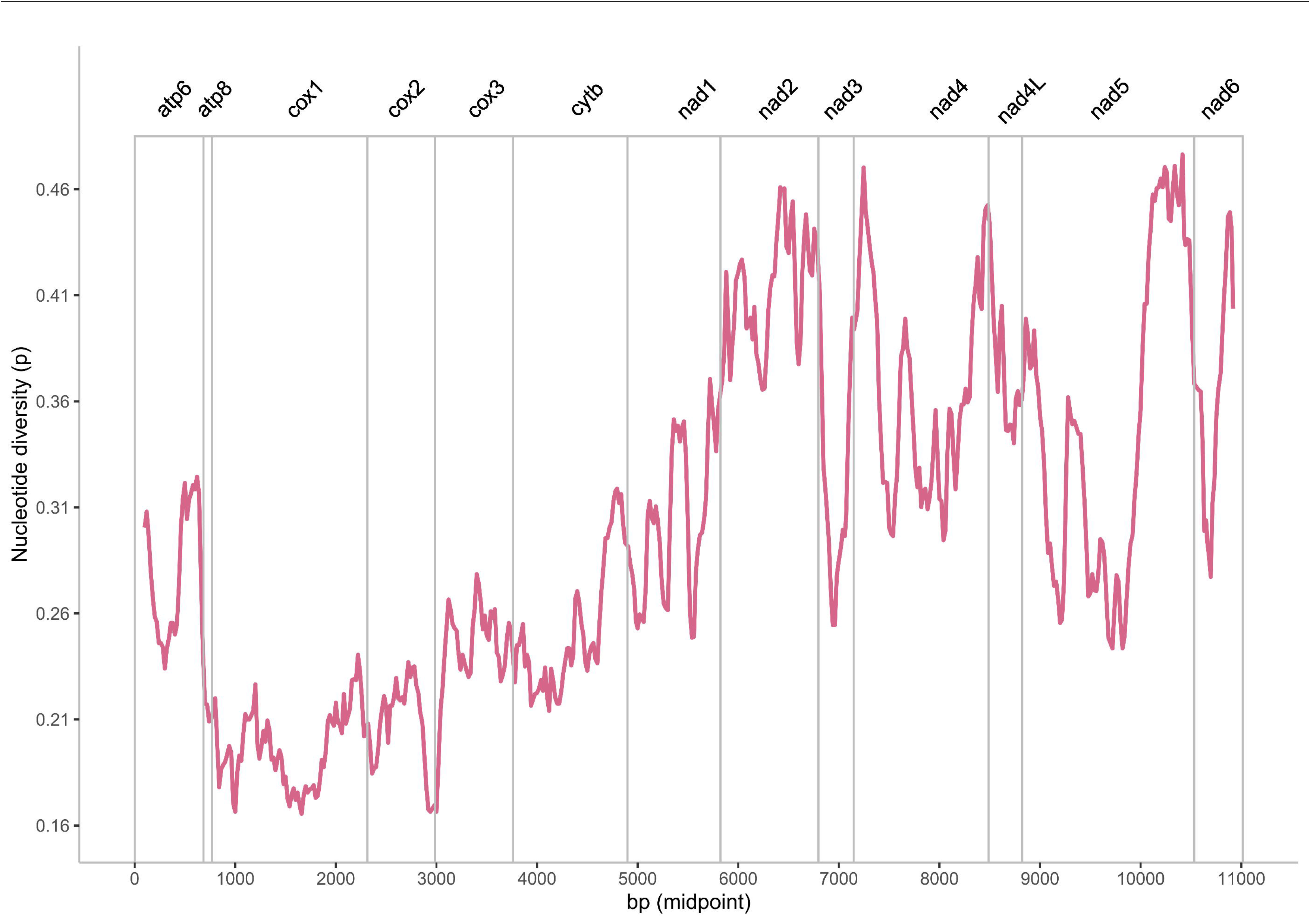
Sliding window analysis of the alignment across 13 protein coding regions among ergasilid copepods (*Ergasilus kandti, E. tumidus, Sinergasilus major, S. polycolpus* and *S. undulatus*). The line shows the value of nucleotide diversity (π) with a window size of 200 bp and a step size of 20; the value is inserted at its mid-point.

**Fig. 6:**
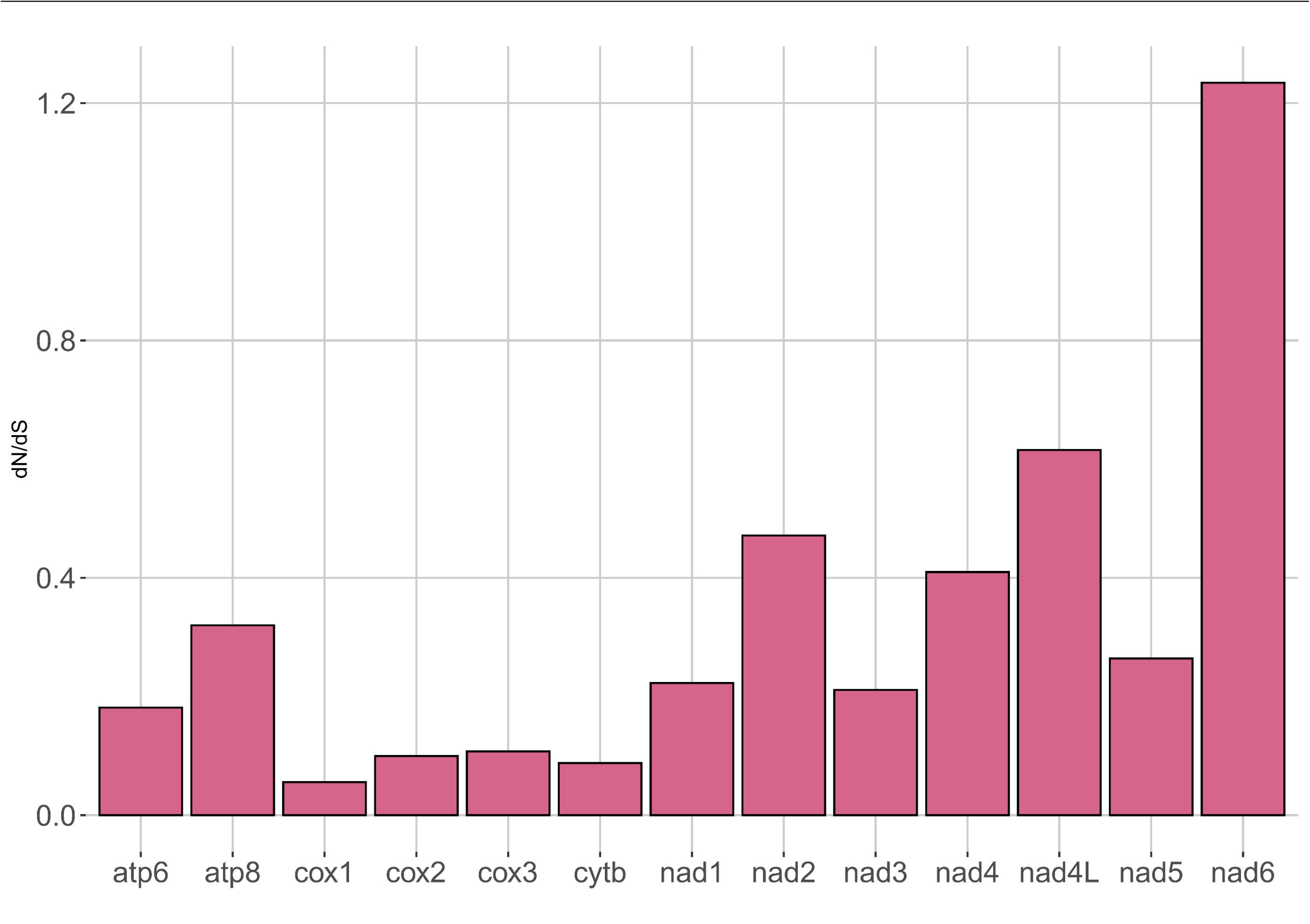
Graphical representation of the ratios between the total number of non-synonymous (dN) and synonymous (dS) nucleotide substitutions per mitochondrial protein coding region across ergasilid copepods (*Ergasilus kandti, E. tumidus, Sinergasilus major, S. polycolpus*, and *S. undulatus*).

### Phylogenetic relationships

We applied maximum likelihood (ML), Bayesian inference (BI), and maximum parsimony (MP) analyses to a concatenated alignment of partial 18S rDNA and 28S rDNA sequences of ergasilids (1500 bp, 212 informative sites). These different analyses resulted in phylogenetic trees with congruent topologies. Therefore, only the ML tree is shown in Fig. 7. The resulting tree topology supports four clades within Ergasilidae including the three clades (Clade I–III) reported in previous studies (see Song et al., 2008). Furthermore, a fourth clade (Clade IV) includes all sequenced African species of *Ergasilus*, namely *E. caparti, E. parvus, E. parasarsi, E. macrodactylus*, *E. mirabilis*, *E. megacheir*, and *E. kandti* as well as the Malagasy *Dermoergasilus madagascarensis* Míč, Řehulková, Šimková, Razanabolana & Seifertová, 2024 and the non-African species *E. sieboldi* and *E. yaluzangbus* Kuang & Qian, 1985. Clade IV is subdivided into two well-supported groups: the African species of *Ergasilus* and the Asian species *E. yaluzangbus* on one hand and *D. madagascarensis* and *E. sieboldi* on the other. Our results also indicate that *Ergasilus* is rendered paraphyletic by *Dermoergasilus* Ho & Do, 1982, *Neoergasilus* Yin, 1956, *Paraergasilus* Markevich, 1937, and *Sinergasilus* Yin, 1949. Phylogenetic analyses (ML, BI, MP) of *cox1* deviated from the rDNA-based topology (Supplementary Fig. S3), e.g. the myicolid and mytilicolid copepods appear to be nested inside Ergasilidae and the monophyly of *Sinergasilus* has low support. Generally, support values were lower throughout the *cox1* phylogeny, leading to many deeper nodes being collapsed into polytomies.

**Fig. 7:**
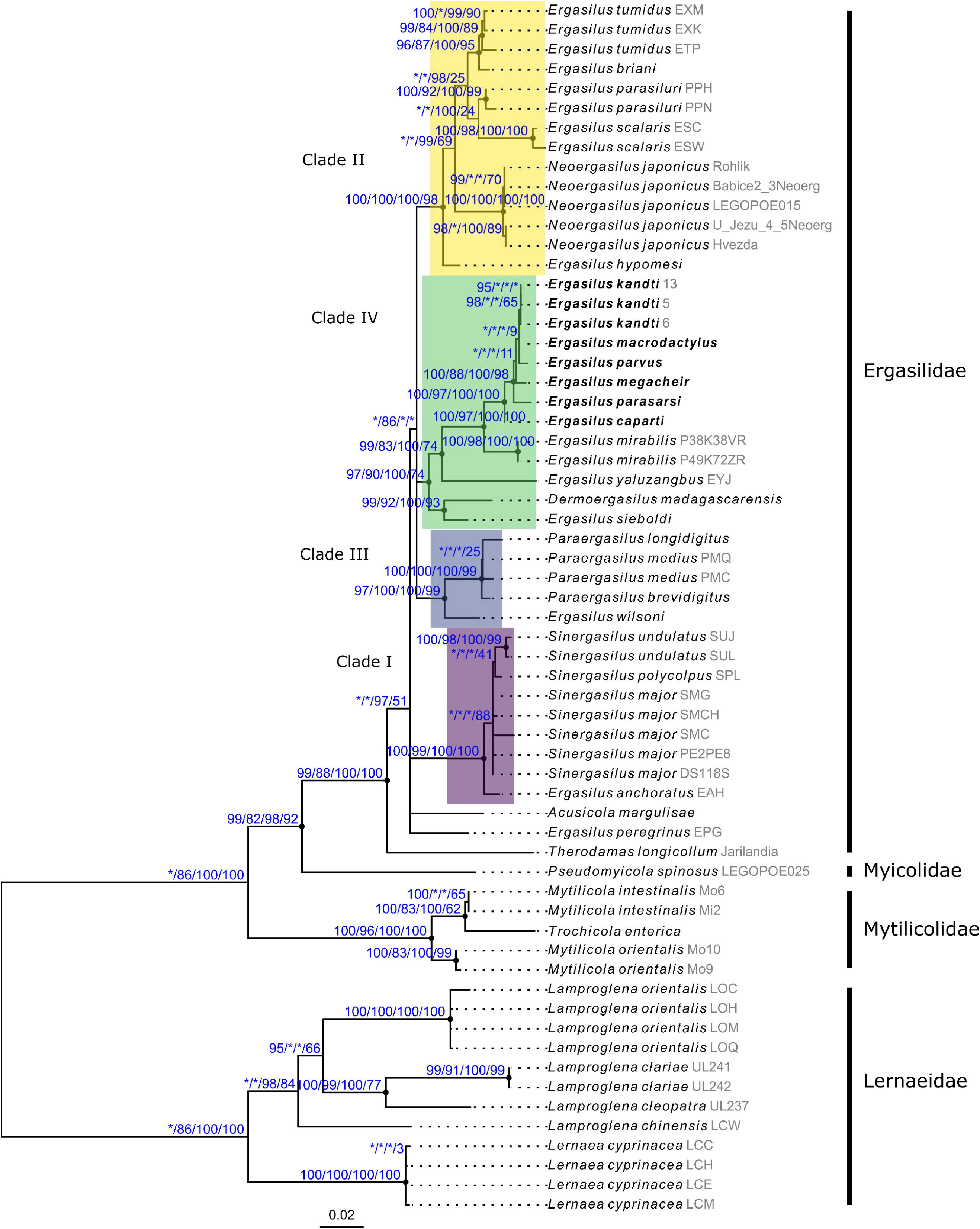
Maximum-likelihood phylogram of ergasilid copepods based on partial 18S and 28S rDNA sequences also including mytilicolid, myicolid, and lernaeid specimens as outgroups (used for rooting). Support is indicated at nodes with ultrafast bootstrap values (UFBoot) and Shimodaira–Hasegawa-like approximate likelihood ratios (SH-aLRT), followed by Bayesian posterior probabilities (PP), and GC values inferred through symmetric resampling (PD=D0.33) in parsimony analysis; asterisk (*) indicates low or moderate support below the threshold (UFBootD<D95, SH-aLRTD<D80, PPD<D95, and GC < 0); black dots, internal nodes with strong support across all support values; labels in bold are species occurring in Lake Tanganyika; labels in grey refer to isolate numbers in Table 1. Clade numbering (I-III) follows Song et al. (2008), with Clade IV added in the present study. Scale bar: estimated number of substitutions per site.

## Discussion

### Morphology and host repertoire of *E. kandti*

We investigated parasitic copepods infecting a specimen of *Tylochromis polylepis* from Lake Tanganyika in East Africa, one of the world’s biodiversity hotspots, and with a so far understudied copepod diversity. A morphological examination based on light and confocal laser microscopy revealed that *T. polylepis* was infected by *Ergasilus kandti* and morphological features of the parasites largely matched those listed in previous characterizations (van Douwe, 1912; Capart, 1944). However, we failed to observe the variation of short setae associated with the caudal rami reported by Capart (1944). Our samples only presented two of these setae as reported by the original description (van Douwe, 1912).

Previous studies on ergasilid copepods have mostly applied light microscopy to highlight systematically informative characters. In recent decades, other imaging techniques have found their way into ergasilid research. For instance, scanning electron microscopy was employed to visualize the pathological alterations in host gill filaments and the feeding apparatus (Kilian & AvenantLOldewage, 2013; Dos Santos et al., 2021; Avenant-Oldewage et al., 2024) as well as systematically informative characters on the legs (Murray et al., 2016). Confocal laser microscopy has been applied successfully to caligids, tisbids, calanids, centropagids, cyclopids, and peltidiids (Buttino et al., 2003; Michels, 2007; Zirbel et al., 2007; Fitzer et al., 2012; Kaji et al., 2012), but this technique has remained absent from the ergasilid literature. Here, we applied confocal laser microscopy to ergasilid copepods for the first time using Congo Red as a fluorescent marker for the exoskeleton. Confocal laser microscopy revealed no additional features compared to traditional light microscopy and was more time-intensive. However, it provides a three-dimensional (i.e. more natural) view of the organism and requires no clearing agent. Therefore, we suggest that this technique adds spatial context to characters that require flattening to be observed in a single plane through light microscopy.

Lake Tanganyika contains several examples of adaptive radiations, i.e. explosive speciation, including snails, fishes (Salzburger et al., 2014), and flatworms (Vanhove et al., 2015; Brand, 2023). The best-known examples are cichlid fishes with around 250 species (Ronco et al., 2020). However, *T. polylepis*, the host of *E. kandti*, is only a recent colonizer of Lake Tanganyika (Koch et al., 2007). The parasites of *T. polylepis* are, hence, of particular interest in terms of studying the effects of adaptive radiations on symbiotic relationships. Prior studies suggest that ergasilid copepods often infect a broad array of host species, e.g., *E. sieboldi* is reported from 15 fish families (Walter & Boxshall, 2023). *Ergasilus kandti* appears to follow this trend. The species has previously been reported from Lake Tanganyika parasitizing both the cichlid radiation flock (Fryer, 1965) as well as a late arrival [see also *E. kandti* infecting *Oreochromis tanganicae* (Fryer, 1965)]. In contrast, no monogenean flatworms shared between *T. polylepis* and other cichlid species in the lake were reported so far (Muterezi Bukinga et al., 2012; Rahmouni et al., 2018; CruzLLaufer et al., 2022). The distribution range of *E. kandti* is not limited to Lake Tanganyika (Fig. 1). This wide geographical range and host repertoire (Fig. 1, Supplementary Table S1) aligns with the low environmental specificity reported in other ergasilid copepods (Mathews et al., 2017; Fikiye et al., 2023).

### Ergasilidae: clades and nomenclature

Due to the persistent lack of DNA sequence data, inferring any biogeographical patterns in the evolution of ergasilid copepods has long been a challenge. As noted by Míč et al. (2023) and mentioned above, these data have a pronounced geographical bias as all molecular data until 2019 were from specimens collected in Asia. The phylogenetic relationships of ergasilid copepods remain generally poorly understood as only 11% (31, see Table 1) of described ergasilid species [275, see Walter & Boxshall (2023)] have been sequenced to date. Here, the phylogenetic analyses suggest that ergasilid species reported from Lake Tanganyika including *E. kandti* form a monophyletic group (part of Clade IV, Fig. 7). No DNA sequences of other African species have been published, bar *E. mirabilis* (Fikiye et al., 2023) as well as the recently described *D. madagascarensis*. The latter species is reported from Madagascar (Míč et al., 2024) and is more closely related to *E. sieboldi*, a species from Eurasia, suggesting that at least two lineages of ergasilids are present in Africa. Nonetheless, low taxonomic coverage alongside the high evolutionary rates of parasitic organisms (Bromham et al., 2013; Vanhove et al., 2013; Gao et al., 2022) might also explain the difficulty of resolving the deeper nodes of the ergasilid phylogeny, especially in the case of the *cox1* phylogeny, which involves sequences of fewer species than the ribosomal phylogeny.

From a systematic point of view, morphological characters widely used to distinguish lineages of Ergasilidae appear to result in mixed success. Several genera, each with their own set of diagnostic characters, including *Neoergasilus*, *Paraergasilus*, and *Sinergasilus*, are repeatedly confirmed as monophyletic groups (Song et al., 2008; Santacruz et al., 2020; Kvach et al., 2021; Míč et al., 2023, 2024). At the same time, these genera render *Ergasilus* a paraphyletic taxon as suggested in previous studies (Song et al., 2008; Santacruz et al., 2020; Kvach et al., 2021; Míč et al., 2023, 2024). Some species including *E. anchoratus* Markevich, 1946 and *E. wilsoni* Markevich, 1933 appear as sister taxa to *Sinergasilus* and *Paraergasilus* (Fig. 7), yet they were not placed in these genera by the authors erecting these genera, respectively. The nomenclatural history of the ergasilids further highlights the difficulty of creating generic diagnoses that allow to effectively distinguish groups. *Pseudergasilus* Yamaguti, 1936, *Trigasilus* Fryer, 1956, and *Ergasiloides* Sars, 1909 are three genera that were synonymized, *Pseudergasilus* and *Ergasiloides* with *Ergasilus*, and *Trigasilus* with *Paraergasilus*, after authors of subsequent publications found the diagnoses to be inadequate. *Pseudergasilus* was differentiated by having “thoracic segments that are distinctly segmented from each other” (Yamaguti, 1936), but Kim & Choi (2003) suggested that this difference was merely an artifact of the growth state of the copepods. *Trigasilus* was created by Fryer (1956). The author later admitted to not being aware of the original diagnosis of *Paraergasilus* (Markewitsch, 1937) but also remarked on the difference in the attachment sites (gill chamber *vs.* gills) and it should be noted that *Paraergasilus minutus* (Fryer, 1956) (synonym: *Trigasilus minutus*) is the only African freshwater representative of this genus (but see *Paraergasilus lagoonaris* Paperna, 1969, which is reported from brackish water fishes in Ghana).

Remarkable in the context of the present study is *Ergasiloides.* This genus was created to encompass three species from Lake Tanganyika (*E. megacheir*, *E. macrodactylus*, and *E. brevimanus*), the two former of which form part of the Lake Tanganyika group in the present study (highlighted in bold as part of Clade IV, Fig. 4). The genus was synonymized with *Ergasilus* because Sars (1909) based his diagnosis on immature specimens according to Fryer (1965). However, Fryer (1965) also noted that species considered as members of *Ergasiloides* by Sars (1909) differed from other species of *Ergasilus* because adult females have two instead of three abdominal somites (or ‘urosomites’) but opted to emend the generic diagnosis of *Ergasilus* to incorporate both this feature as well as the five-segmented antennule of *E. kandti* as he considered these characters mere ‘anomalies’ and variations to be expected for this group of parasites. The abdominal somites are indeed not fused in several species from Lake Tanganyika, including *E. kandti* (van Douwe, 1912), *E. parasarsi*, and *E. parvus* (Míč et al., 2023), and a recent re-description also refutes this feature in *E. macrodactylus* (Míč et al., 2023). Nonetheless, our phylogenetic analysis (Fig. 7) suggests that if the genus *Ergasiloides* were to be re-established, it could be applied as the name of the Lake Tanganyika group (part of Clade IV) as it was previously assigned to several of these species. However, this step would require the provision of a consistent generic diagnosis through a systematic revision.

### Mitochondrial genomes of ergasilids: tRNA rearrangement and nucleotide polymorphism

In general, crustaceans have high rates of gene order rearrangement in mitochondrial genomes (Sterling-Montealegre & Prada, 2024). Given the general scarcity of ergasilid DNA sequences, it comes as no surprise that full mitochondrial genomes are available for even fewer species. Nonetheless, a comparison of ergasilids and other copepods can provide some insights into the interspecific variation of these sequences and the gene order. While the order of mitochondrial rDNA and rRNA regions is stable across ergasilid copepods, we detected several instances of rearrangement between different tRNAs in the mitochondrial genome. Furthermore, we report the presence of the TA-abbreviated stop codon (15.38%) in the protein coding genes being higher than the average usage (2.01%) reported mitochondrial regions at the level of Copepoda (He et al., 2023). In comparison to free-living copepods (Ki et al., 2009; Minxiao et al., 2011), ergasilid copepods show the complete absence of repetitive regions and a relatively short non-coding region in their mitochondria (in those assembled so far).

We also observed similar patterns found in other parasites when comparing nucleotide polymorphism and dN/dS ratios. Nucleotide polymorphism across the mitochondrial rrDNA regions of ergasilid copepods was disproportionately higher in regions coding for dehydrogenase subunits compared to cytochrome and ATP coding regions (see Figs. 5 and 6). Our results on the dN/dS ratio further suggest that the *cox1* gene is under the strongest selection pressure. This pattern aligns with the results of previous studies highlighting low evolutionary rates of the *cox1* compared to other mitochondrial protein coding regions, e.g., in calanoids (Zhang et al., 2021), but also other copepods (He et al., 2023; Huang et al 2024). Similar patterns are also present in monogenean flatworms infecting cichlid (Kmentová et al., 2021) and clupeid fishes (Kmentová et al., 2023) in Lake Tanganyika. These parasites share not only the gill habitat and host lineages (i.e. cichlids) with ergasilid copepods, but also both have a single-host life cycle, However, the limited number of ergasilid copepods of which the mitochondrial genomes has been assembled today (n = 5, including the present study) in comparison to the estimated worldwide diversity of 275 species prevents any general conclusion and more studies are needed to unravel (mito-)genomic patterns underlying diversification processes of ergasilid copepods.

## Conclusions

The East African lakes and, specifically Lake Tanganyika, are a biodiversity hotspot for many different vertebrate and invertebrate taxa. Despite this knowledge, copepods—specifically parasitic species—are one of the most overlooked groups in these lakes. A major knowledge gap also remains associated with the lack of molecular data across the geographical ranges of ergasilid species. We suggest that to understand the drivers of evolution of ergasilid copepods, these parasites need to be sampled across their geographical range. Specifically, *E. kandti* should be sampled in different water bodies and, specifically, hosts inside and outside of Lake Tanganyika. Furthermore, ergasilid copepods were suggested as bioindicators being sensitive to eutrophication and other processes derived from ecosystem degradation (Sasal et al., 2007). Therefore, studying temporal dynamics of ergasilid copepods, species combining both free-living and parasitic life stages, could be informative for ecosystem health assessment.

Our study provides an insight of how upscaling both morphological characterization and molecular analyses using technological solutions applied for other copepods and metazoan parasites can expand our knowledge on this key taxon. While the use of next-generation sequencing and confocal imaging is new to ergasilids and other copepods in East Africa, protocols can be transferred without facing major challenges. We encourage researchers to incorporate these tools into future studies on ergasilids worldwide.

## Data availability

Parasite voucher material was deposited in the collection of Hasselt University under accession numbers HU XXIII.1.41–HU XXIII.1.50, HU XXIII.2.01, and HU T-I.1–HU T-I.11. The ribosomal DNA sequences were uploaded to GenBank and can be found under accession numbers PQ249839–PQ249843. The mitogenome DNA sequence is available in the GenBank Nucleotide Database under the accession number PQ276880. Raw Illumina reads were submitted to SRA (accession number: SRR30471034). All sequences are linked under BioProject accession PRJNA1153390. Image files and phylogenetic tree files can be accessed at MorphoBank (http://www.morphobank.org/, Project 5467).

## Acknowledgements

We would like to dedicate this manuscript to Geoffrey Fryer (1927–2024), whose works proved essential for the present study. We would like to thank Stephan Koblmüller, Holger Zimmermann, Simona Georgieva, Gyrhaiss Kapepula Kasembele, Cyprian Katongo and Taylor Banda, for their help in organising and conducting field work. We further thank Natascha Steffanie and Martijn Heleven for assistance concerning laboratory work. We would also like to thank the anonymous reviewers for the valuable feedback on this manuscript.

## Author’s contribution

DJ, NK, and AJCL conceptualized the study. NK led the sampling campaign with the assistance of JV and LM. LM collected the fish host specimen in the field. AJCL collected parasite specimens off the host gills. DJ prepared samples and conducted imaging and molecular analyses under supervision of NK and AJCL. NK and AJCL analyzed next-generation sequencing data. DJ and AJCL performed phylogenetic analyses. DJ, NK, and AJCL wrote the article with input from MPMV, JV and LM. All co-authors approved the final version of the manuscript.

## Ethical standards

Fieldwork was carried out with the approval of the competent local authorities under the permission of the Fisheries Department of Zambia and under a study permit issued by the government of Zambia (SP 008732).

## List of supplementary material

**Supplementary Table S1**. Recorded locations of *Ergasilus kandti*. Geographical coordinates were estimated based on reports in the literature (i.e. geographical centers of lakes, approximate locations indicated in figures etc.)

**Supplementary Table S2**. Counts of spines and setae of specimens of *Ergasilus kandti* collected in the present study. Empty cells indicate missing (unobserved) data. Cells highlighted in yellow indicate deviations from the spine-seta formula of the species.

**Supplementary Fig. S3**. Maximum-likelihood phylogram of ergasilids copepods based on partial *cox1* sequences. Support is indicated at nodes with ultrafast bootstrap values (UFBoot) and Shimodaira–Hasegawa-like approximate likelihood ratios (SH-aLRT), followed by Bayesian posterior probabilities (PP), and GC values inferred through symmetric resampling (P□=□0.33) in parsimony analysis; asterisk (*) indicates low or moderate support below the threshold (UFBoot□<□95, SH-aLRT□<□80, PP□<□95, and GC < 0); black dots, internal nodes with strong support across all support values. Labels in grey refer to isolate numbers in Table 1. Scale bar: estimated number of substitutions per site.

