## Supplementary Table S1 for "Mitogenomics, phylogenetic position, and updated distribution of *Ergasilus kandti*, an ergasilid copepod parasitizing African cichlid fishes"

**Supplementary Table S1**. Recorded locations of *Ergasilus kandti*. Geographical coordinates were estimated based on reports in the literature (i.e. geographical centers of lakes, approximate locations indicated in figures etc.)

| **Host** | **Location** | **LAT** | **LON** | **Reference** | **Remarks** |
| --- | --- | --- | --- | --- | --- |
| *Citharinus citharus* | Lake Volta, Black and White Volta confluence | 7.626 | 0.109 | Paperna (1969) |  |
| *Hemisynodontis* sp*.* | Lake Volta, Black and White Volta confluence | 7.626 | 0.109 | Paperna (1969) |  |
| *Lates niloticus* | Lake Volta, Black and White Volta confluence | 7.626 | 0.109 | Paperna (1969) |  |
| *Synodontis membranaceus* | Lake Volta, Black and White Volta confluence | 7.626 | 0.109 | Paperna (1969) |  |
| *L. niloticus* | Gourao, Niger Basin | 13.133 | 2.317 | Capart (1956) |  |
| *Bagrus bajad* | Lake Albert | 1.819 | 31.326 | Thurston (1970) |  |
| in plankton | Lake Albert | 1.683 | 30.917 | van Douwe (1912) | geographical center |
| *L. niloticus* | Lake Albert | 1.819 | 31.326 | Fryer (1965), Thurston 1970 |  |
| *Lamprologus lemairii* | Lake Tanganyika | -6.100 | 29.500 | Fryer (1965) | geographical center |
| *Limnotilapia dardennii* | Lake Tanganyika | -6.100 | 29.500 | Fryer (1965) | geographical center |
| *Oreochromis tanganicae* | Lake Tanganyika | -6.100 | 29.500 | Fryer (1965) | geographical center |
| *Plecodus paradoxus* | Lake Tanganyika | -6.100 | 29.500 | Fryer (1965) | geographical center |
| *Pseudosimochromis curvifrons* | Lake Tanganyika | -6.100 | 29.500 | Capart (1944) | geographical center |
| *Tilapia* sp. | Lake Tanganyika | -6.100 | 29.500 | Fryer (1965) | geographical center |
| *Tylochromis polylepis* | Lake Tanganyika | -8.767 | 31.117 | this study |  |
| *Pterochromis congicus* | Lake Tumba | -0.833 | 18.000 | Fryer (1964) |  |
| *L. niloticus* | Lower Nile | 9.372 | 31.547 | Fryer (1968) | points on map |
| *Tylochromis bangwelensis* | Luapula River | -10.367 | 28.633 | Fryer (1967) | RMCA museum records |
| *Tylochromis myolodon* | Luapula River, N'Kole | -9.430 | 28.530 | Fryer (1967) |  |
| *T. polylepis* | Malagarasi Delta | -5.233 | 29.783 | Fryer (1967) | erroneously placed in Rwanda |
| *L. niloticus* | River Niger | 13.959 | -5.366 | Fryer (1968) | points on map |
| *T. mylodon* | Lake Mweru | -8.730 | 28.691 | Fryer (1967) |  |
