## Supplementary figures and images for "Mitogenomics, phylogenetic position, and updated distribution of *Ergasilus kandti*, an ergasilid copepod parasitizing African cichlid fishes"

### SupplementaryFigS3.png

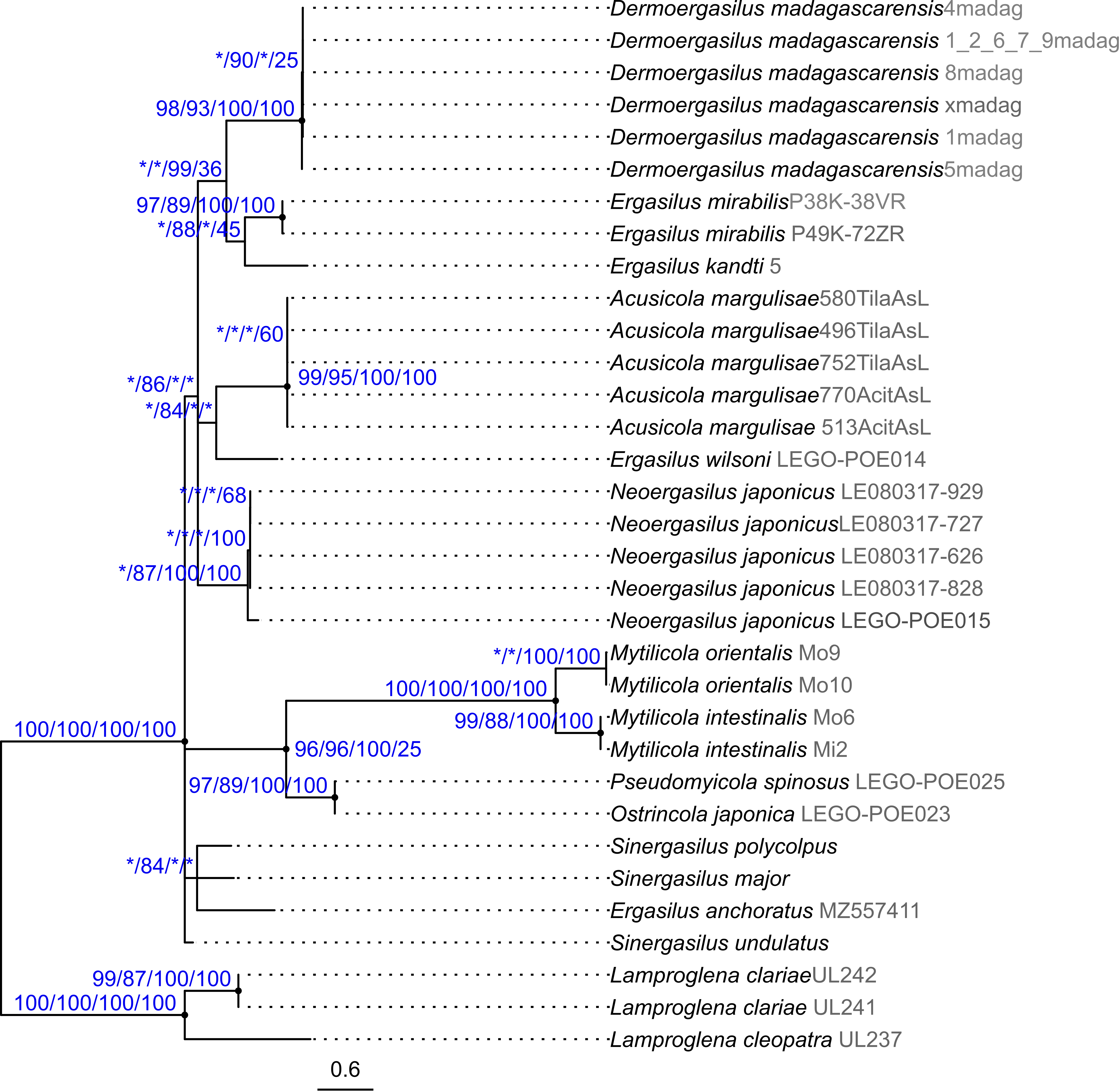
